# Intercellular Signaling Network Underlies Biological Time Across Multiple Temporal Scales

**DOI:** 10.1101/315788

**Authors:** Joshua Millstein, Keith C. Summa, Xia Yang, Jun Zhu, Huaiyu Mi, Martha H. Vitaterna, Fred W. Turek, Bin Zhang

## Abstract

**Motivation:** Cellular, physiological and molecular processes must be organized and regulated across multiple time domains throughout the lifespan of an organism. The technological revolution in molecular biology has led to the identification of numerous genes implicated in the regulation of diverse temporal biological processes. However, it is natural to question whether there is an underlying regulatory network governing multiple timescales simultaneously.

**Results:** Using queries of relevant databases and literature searches, a single dense multiscale temporal regulatory network was identified involving core sets of genes that regulate circadian, cell cycle, and aging processes. The network was highly enriched for genes involved in signal transduction (P = 1.82e-82), with p53 and its regulators such as p300 and CREB binding protein forming key hubs, but also for genes involved in metabolism (P = 6.07e-127) and cellular response to stress (P = 1.56e-93). These results suggest an intertwined molecular signaling network that affects biological time across multiple temporal scales in response to environmental stimuli and available resources.

**Supplementary information:** Supplementary data are available online.

## 1 Introduction

The passage of time is a fundamental feature of life. Although time is often considered to be linear, organisms must organize developmental, cellular and physiological processes across multiple time domains. For example, segmentation and patterning in animals requires the initiation of precise temporally ordered molecular programs throughout development to ensure correct morphology of the organism. Although the specific length of time that this process takes differs between species, the temporal pattern is largely conserved, indicating universal control mechanisms involved in regulating the timing of development. On a separate time scale, the regular rotation of the Earth about its axis every 24 hours leads to predictable daily changes in the environment. Organisms have evolved endogenous circadian rhythms to anticipate these changes and organize behavior and internal physiology to occur at the correct time in relation to cycles in the external environment. A set of well-conserved circadian clock genes encodes a cell-autonomous pacemaker characterized by a transcriptional-translational feedback loop that is itself modulated and reciprocally regulated by hundreds of other genes, including several key metabolic regulatory genes. The circadian system controls the expression of thousands of genes (i.e., clock-controlled genes) in nearly all cells of the body, establishing overt rhythms in cellular and physiological function at the tissue- and organ-system levels. On yet another time scale, individual cells progress through a cycle of quiescence and division that enables the maintenance of a stable population of cells while allowing for the division, differentiation and replenishment of specific cell types. Cell division and differentiation require numerous complex temporally ordered processes, such as DNA replication, chromatin condensation and chromosome alignment and segregation into daughter cells. All organisms also experience characteristic age-related changes that occur throughout the lifespan and most experience deleterious changes that accumulate over time, eventually resulting in age-related diseases, senescence, and death.

Thus, cellular, physiological and molecular processes must be organized and regulated across multiple time domains throughout the lifespan. The technological revolution in molecular biology has led to the identification of numerous genes implicated in the regulation of circadian rhythms, the cell cycle and aging. We here report a systems-level analysis using various network methodologies to systematically investigate whether constituents of temporal regulation within a single time domain are involved in others as well, that organisms utilize common sets of time regulatory genes and networks to ensure multi-scale temporal organization. Such analysis is expected to reveal insight into the common cellular and molecular mechanisms used in the regulation of timing across multiple domains. Indeed, this concept that genes with a known role in one domain have critical functions in another has support in the literature: cyclin A and its regulator, both essential cell cycle factors, were recently shown to regulate sleep in *Drosophila* via activity in a small set of post-mitotic neurons associated with the circadian clock (Rogulja and Young, 2012). Interestingly, as described below, one of the critical multi-scale nodes (i.e., genes present across the three time scales examined: circadian, cell cycle and aging) revealed in our analysis is cyclin-dependent kinase 6 (CDK6). Given the well-described role of the cyclins and cyclin-dependent kinases in the cell cycle, traditionally it would not be expected to observe such a gene to be active in post-mitotic neurons. However, given our analyses of coordinated temporal regulation across multiple domains using common factors, it is reasonable that specific genes and pathways be examined for important roles related to the regulation of timing across scales.

## 2 Methods

The ultimate objective was to find evidence of an underlying gene network affecting biological time across multiple scales. The overall approach involved identifying sets of genes that regulate temporal processes for several time scales, determining regulatory relationships between these genes, and finding links that operate across time scales. Core components of the resulting multi-scale network would likely affect temporal biological processes generally.

### 2.1 Defining Timekeeping Genes

Core sets of genes, ‘timekeepers’, that regulate human temporal biological processes at the i) circadian, ii) cell cycle, and iii) ageing scales were defined through queries of relevant databases and literature searches. Circadian clock genes were identified using several comprehensive reviews of the genetics and physiology of mammalian circadian rhythms and one genome-wide RNAi screen for genes that modulate circadian rhythms in cultured human cells (Lowrey and Takahashi, 2011; Rosenwasser and Turek, 2011; Vitaterna and Turek, 2011; Zhang, et al., 2009). These resources were supplemented with manual examination of each of the reference lists and PubMed searches using the MeSH subject headings “circadian rhythms” and “genes”. In addition, the Molecular Signatures Database (MSigDB) (Subramanian, et al., 2005) was used to download REACTOME circadian clock pathway genes, KEGG, circadian rhythm genes from the Kyoto Encyclopedia of Genes and Genomies (KEGG) (Kanehisa, et al., 2016), and circadian pathway genes from the Pathways Intereaction Database (PID). The Human Ageing Genomic Resources (HAGR) (Tacutu, et al., 2013) was used to identify genes involved in regulation of human aging processes. Genes involved in human longevity were downloaded from MSigDB, curated by BioCarta. Genes involved in regulating the cell cycle were also identified from MSigDB sets, REGULATION OF CELL CYCLE, KEGG CELL CYCLE, and REACTOME CELL CYCLE MITOTIC.

### 2.2 Reconstructing a Multiscale Network of Timekeepers

There are many types of dependencies that can define gene-gene relationships as well as a diverse array of methods for identifying them. Our intent was to determine whether a core regulatory network that operates across temporal scales exists, and if so to reconstruct it. The approach involves using a variety of public resources and databases to identify links with strong evidence of dependence. The following methods were used:

#### Coexpression

We made use of a compendium of previously reconstructed gene coexpression networks from multiple human cohort studies (Chen, et al., 2008; Emilsson, et al., 2008; Lum, et al., 2006; Tran, et al., 2011; Wang, et al., 2012; Yang, et al., 2010). Gene co-expression network analysis (GCENA) has been increasingly used to identify gene subnetworks for prioritizing gene targets associated with a variety of common human diseases such as cancer and obesity (Chen, et al., 2008; Emilsson, et al., 2008; Gargalovic, et al., 2006; Horvath, et al., 2006). In a gene coexpression network, the nodes represent genes and edges (links) between any two nodes indicate a relationship (a similar expression pattern) between the two corresponding genes. One important end product of GCENA is gene modules comprised of highly interconnected sets of genes. It has been demonstrated that these types of modules are generally enriched for known biological pathways, for genes that associate with disease traits, and for genes that are linked to common genetic loci (Liu, et al., 2015; Schadt, et al., 2008; Werling and Sanders, 2016; Zhang and Horvath, 2005; Zhu, et al., 2008).

#### Bayesian Network

While coexpression networks can provide a global view of how genes coordinate as groups, they can’t explicitly infer causal relationships among genes, which are critical to identify key regulators. Probabilistic causal networks are one way to model such relationships, where causality in this context reflects a probabilistic belief that one node in the network affects the behavior of another either directly or indirectly. Bayesian networks (Zhu, et al., 2004; Zhu, et al., 2012; Zhu, et al., 2007; Zhu, et al., 2008) are one type of probabilistic causal networks that provide a natural framework for integrating highly dissimilar types of data. Unlike co-expression networks, which allow one to look at the overall gene-gene correlation structure at a high level, Bayesian networks (BN) are sparser but allow a more granular look at the relationships and directional predictions among genes or between genes and other traits such as disease. Thus, we constructed a network for each dataset independently. Each Bayesian network was tissue-specific and was constructed using genetic and gene expression data generated from multiple human populations (Zhu, et al., 2004; Zhu, et al., 2007; Zhu, et al., 2008). For each Bayesian Network, a Markov Chain Monte Carlo (MCMC) approach was used to generate thousands of different plausible networks that were then combined by taking the union to obtain a consensus network (Zhu, et al., 2007).

#### PPI

A variety of databases have been developed to catalog protein-protein interactions, experimentally validated in the lab using a variety of assays. The STRING project (Szklarczyk, et al., 2015) has aggregated this information in their “experiments” type of evidence, which is either direct or indirect. The strength of evidence is quantified according to a numeric score, scaled from 0 to 1, where 1 represents complete confidence in the existence of the interaction. We used the stringent threshold of 0.9 to conclude that there was sufficient evidence of a protein-protein interaction corresponding to a pair of genes.

#### Pathway

Another type of evidence aggregated by STRING is pathway knowledge derived from manually curated databases. The stringent confidence threshold of 0.9 was also used for this type.

#### Consensus

To further the objective of identifying a core multiscale temporal regulatory network, edges that were found using more than one of the preceding four methods were combined into a final summary network, robust to the approaches used to determine interactions.

To insure that the identified networks operated across multiple time scales, edges were included only if the two nodes represented more than one time scale (Fig. 1). For instance, an edge that linked two genes involved in cell cycle but not circadian or aging would be excluded, but if at least one of these nodes was involved in aging or circadian processes, the edge would be included. Another example, if an edge linked one node involved in aging and another involved in circadian, it would be included. Thus, the networks would be fragmented and sparse if there were no underlying multiscale temporal network but more highly connected if one exists.

**Figure 1.**
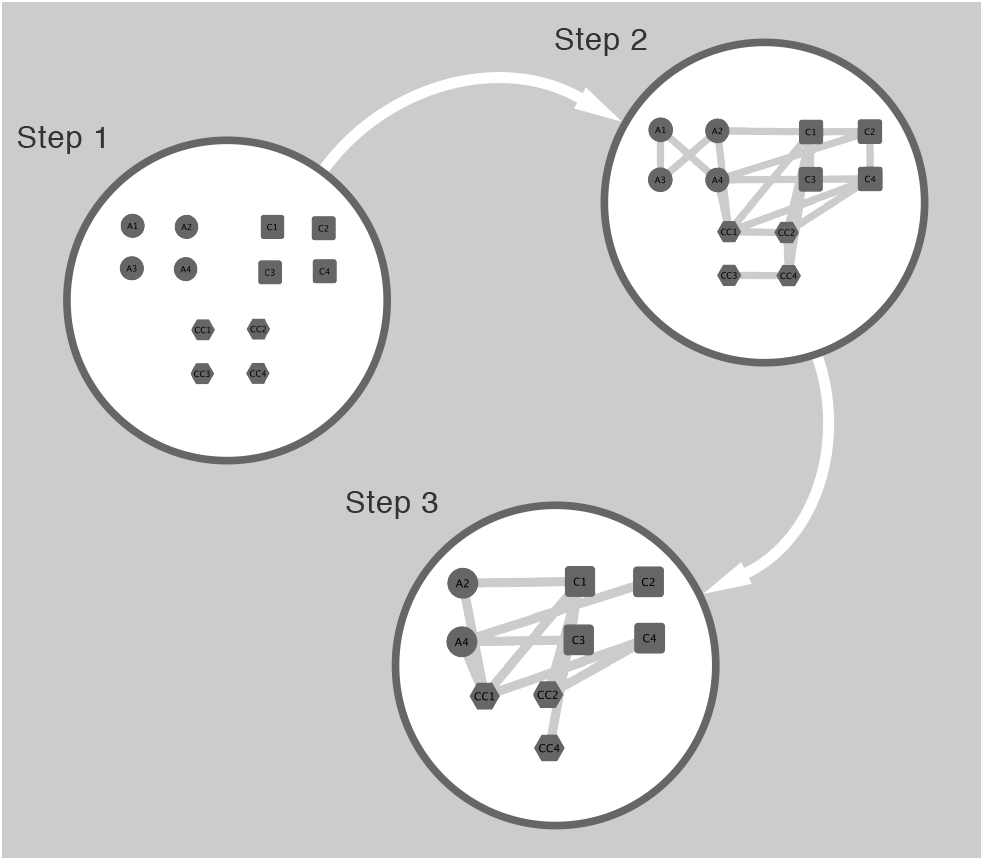
The procedure to construct a multiscale temporal regulatory network includes three steps, 1) identify nodes for multiple temporal scales, 2) determine which nodes are functionally linked, and 3) remove edges that do not span time scales and nodes that do not represent multiple time scales when including first degree neighbors.

### 2.3 Key Driver Analysis

One primary goal of gene network analysis is to identify key regulatory components, or key driver nodes of sub-networks with respect to varying biological contexts. The key drivers will represent the core nodes of the network and tend to have high ‘node degree’, number of edges directly connected to the node. We identified candidate key drivers for each of the subnetworks (evidence types) mentioned. The algorithm takes as input nodes and edges from a network. We first compute the size of the h-layer neighborhood (HLN) for each node. The range of h is from 1 to the diameter of the network. Specifically, for a given node *g*, the size of its HLN is the number of its downstream nodes that are within h edges of *g*. Let ***u*** be an array of the sizes of HLNs and ***d*** be an array of the out-degrees for all the nodes. The nodes are nominated as candidate drivers if their sizes of their HLN are greater than *μ* + *σ* , where *μ* is the mean of ***u*** and *σ* is the standard deviation of ***u***.

## 3 Results

Fig. 2 shows significant overlaps between the three temporal scales. All the three sets highly significantly overlapped with each other, with the aging and cell cycle the most significant (FET P=7.8e-32, 11.2 fold enrichment) followed by aging and circadian (FET P=1.4e-18, 8.2 fold enrichment) and then cell cycle and circadian (FET P=6.0e-13, 5.1 fold enrichment). Clearly, these processes are not independent of each other. While 91 genes were implicated in at least two time-scales, only six had roles supported for all three (Table 1).

**Figure 2.**
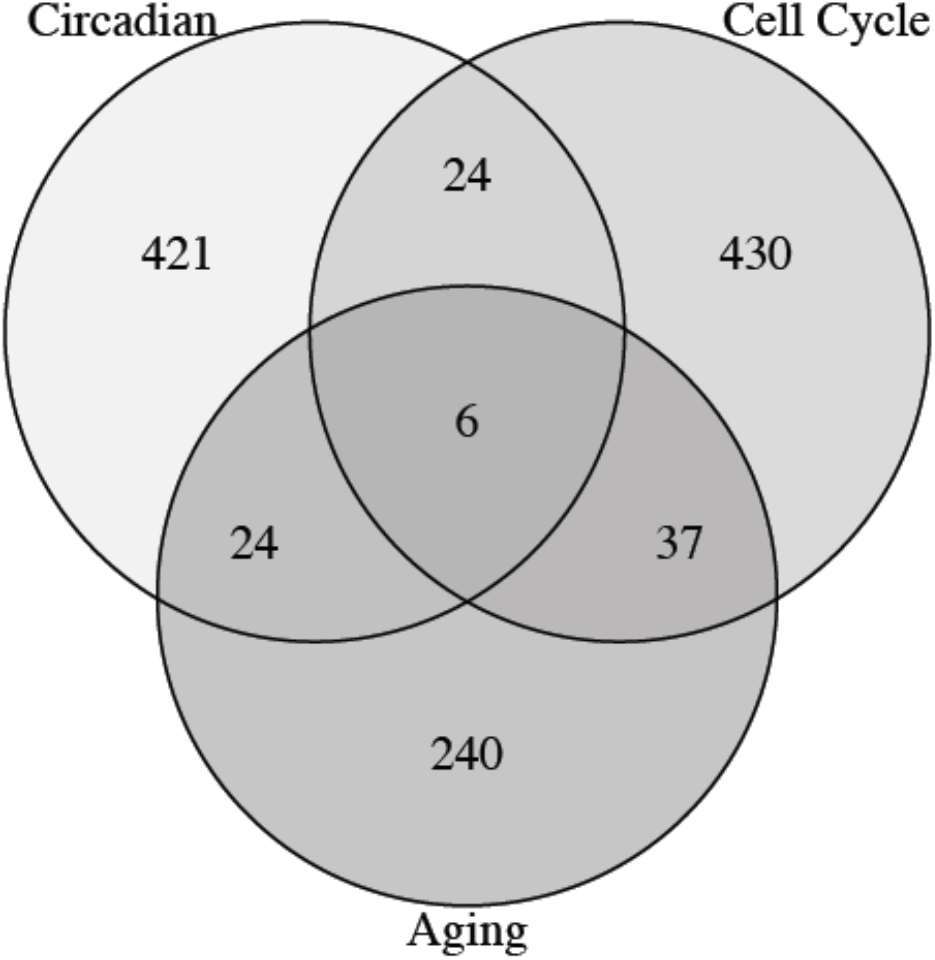
Venn diagram of regulatory genes involved in the temporal scales circadian, aging, and cell cycle.

**Table 1.** Regulatory genes that span circadian, aging, and cell cycle time scales

| Symbol | Name | Source |  |  |
| --- | --- | --- | --- | --- |
|  |  | Circadian | Aging | Cell Cycle |
| *CSNK1E* | Casein Kinase 1 Epsilon | LIT | HAGR | KEGG |
| *GSK3B* | Glycogen Synthase Kinase 3 Beta | LIT | HAGR | KEGG |
| *ABL1* | Abelson Tyrosine-Protein Kinase 1 | LIT | HAGR | KEGG |
| *CREBBP* | CREB Binding Protein | LIT | HAGR | KEGG |
| *EP300* | E1A Binding Protein P300 | LIT | HAGR | KEGG |
| *ATR* | ATR Serine/Threonine Kinase | PID | HAGR | REACTOME |

LIT denotes a search of primary literature. PID, the Pathway Interaction Database [PMID: 18832364]. HAGR, the Human Ageing Genomic Resources [PMID: 23193293]. KEGG, Kyoto Encyclopedia of Genes and Genomes [PMID: 26476454]. REACTOME, curated pathway database [PMID: 26656494].

Three of these, *CSNK1E*, *GSK3B*, and *ATR*, have serine-threonine protein kinase activity, *ABL1*, is a tyrosine kinase, and the remaining two, *CREBBP* and *EP300*, share substantial functional homology and are involved in transcriptional coactivation of many transcription factors. All of these genes play an essential role in cell signaling, the protein kinases by phosphorylating other proteins. *CREBBP* and *EP300* are histone acetyltransferases that play an essential role in multiple signal transduction pathways and are involved in the coordination and integration of signals (Chan and La Thangue, 2001).

### 3.1 Network of Molecular Timekeepers

After identifying interactions between all the timekeepers using STRING, coexpression, and Bayesian network analysis, the resulting network included 39,693 edges (Tables S1 and S2). The large size and high density of the global network necessitated focusing strategies to drill down into biological features and topology of interest. These features include key drivers of the subnetwork of edges that span temporal domains. However, even after filtering out single-domain edges and partitioning by evidence type, the resulting multiscale networks are quite dense, suggesting a complex underlying molecular process that regulates biological time generally, unrestricted to just a single temporal domain. Nodes and network edges are available in supplemental online data.

#### 3.1.1 Multiscale Bayesian Network

The most highly connected region of the Bayesian network (Fig. 3, Table S3) includes the core genes *CENPM*, *MCM5*, *NUSAP1*, *CCNE1 PCNA*, *EXO1*, *ORC1L* and *BUB1B*, which are involved in cell division and more specifically the kinetochore (*CENPM*, *BUB1B*, *NUSAP1*, *CCNE1*) (Li, et al., 2016; Thiru, et al., 2014), connecting microtubules to chromotids. Most of these genes (*CENPM*, *MCM5*, *NUSAP1*, *CCNE1 PCNA*, *ORC1L*, and *BUB1B*), in addition to the key driver BTG2 are targets of P53 (Fischer, et al., 2016; Riley, et al., 2008). Another key driver in this region is HIF1A, a master transcriptional regulator of cellular response to hypoxia that requires recruitment of coactivators such as *CREBBP* and *EP300*. Key drivers *YWHAH* and *PTPN11* are both known to be involved in signal transduction for a variety of cell processes.

**Figure 3.**
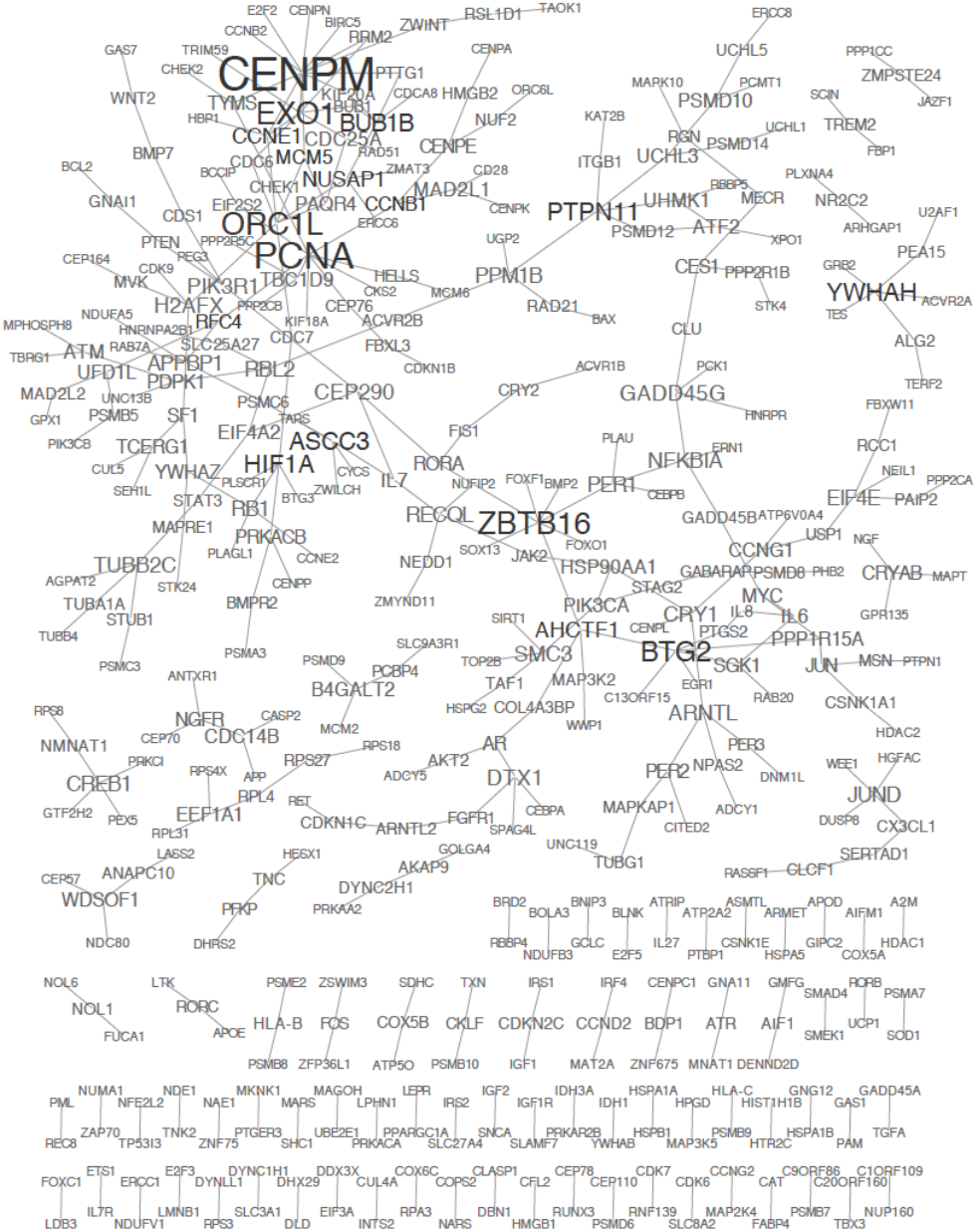
Multiscale Bayesian Network. Nodes are gene symbols of identified molecular timekeepers found to interact with other timekeepers in Bayesian network analyses of genetic and gene expression variation in multiple human cohorts. Edges were included only if they linked two nodes that represented two or more temporal domains. Node size is proportional to node degree. Black indicates key drivers.

#### 3.1.2 Multiscale Coexpression Network

With 1533 edges, this network (Fig. 4, Table S4) was more highly connected than the Bayesian network, which had 371. Key drivers that were also labeled as such in the Bayesian network include *BUB1B*, *PCNA*, and *HIF1A*. As observed above, a substantial proportion of key drivers (*CCNA2*, *BUB1B*, *PCNA*, *FOXM1*, *MAD2L1*, *CENPF*) are regulated by *p53* (*TP53*), also a key driver in this network. Most of the key drivers are involved in cell division and many are also involved the kinetochore (*BUB1B*, *MYC*, *MAD2L1*, *CENPE*, *CENPF*, *TOP2A*) (Li, et al., 2016; Thiru, et al., 2014; Tipton, et al., 2012). *p53* plays a critical role in cellular response to both intrinsic and extrinsic stresses such as DNA damage, which can cause the cell to respond by cell cycle arrest, senescence, or cellular apoptosis (Harris and Levine, 2005). *p53* is in volved in multiple signal transduction pathways and communication of signals to surrounding cells.

**Figure 4.**
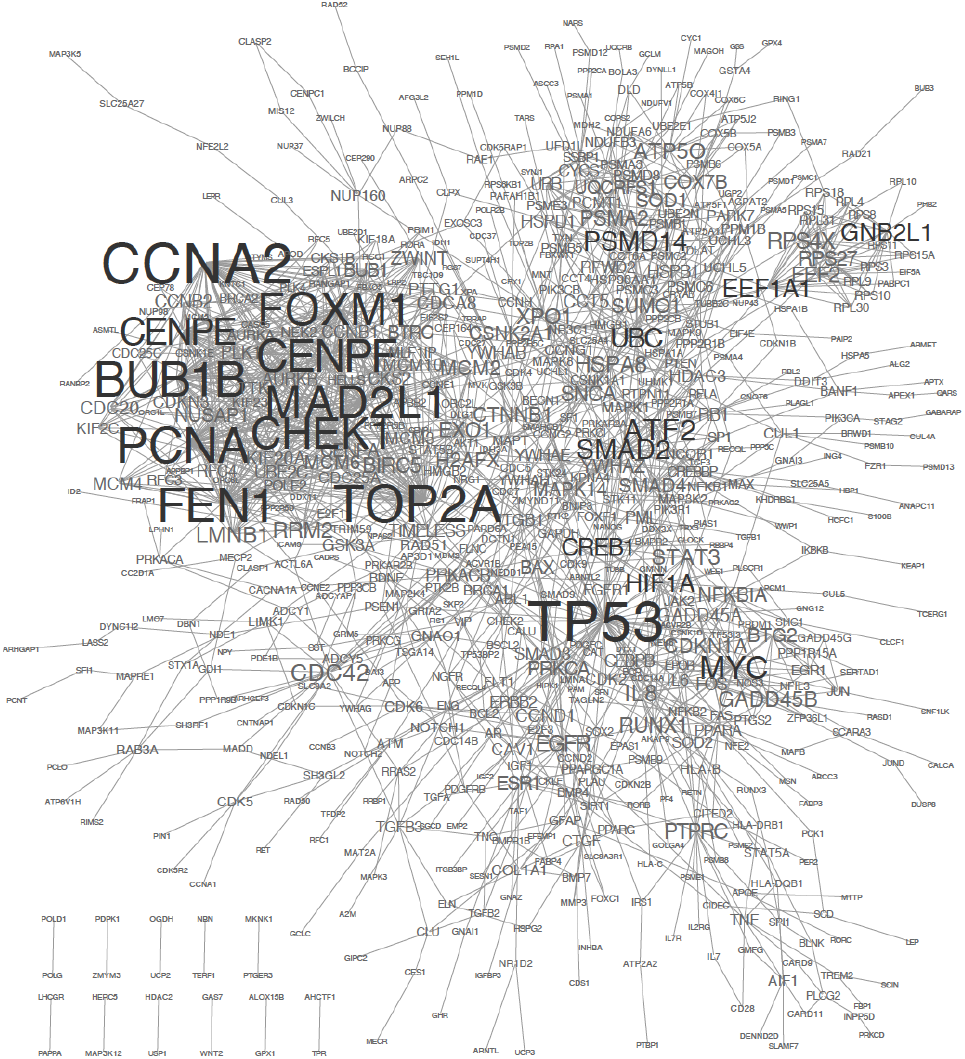
Multiscale Coexpression Network. Nodes are gene symbols of identified molecular timekeepers found to interact with other timekeepers in coexpression network analyses of genetic and gene expression variation in multiple human cohorts. Edges were included only if they linked two nodes that represented two or more temporal domains. Node size is proportional to node degree. Black indicates key drivers.

#### 3.1.3 Multiscale Pathway Network

The pathway network, constructed using the STRING database, was very highly connected with 5446 edges among the 1182 nodes (Fig. 5, Table S5). The key drivers include some reported above, (*TP53*, *CENPM, MAD2L1*, *MYC*, and *BUB1B*). Similarity between the coexpression and PPI networks is also exhibited by interdependence between the key drivers with the highest node degrees, *CCNA2* and *CDK1*. *CCNA2* is essential in mitosis during the G2 to M transition when it activates *CDK1*, which is itself essential for cell cycle progression [Bendris 2011].

**Figure 5.**
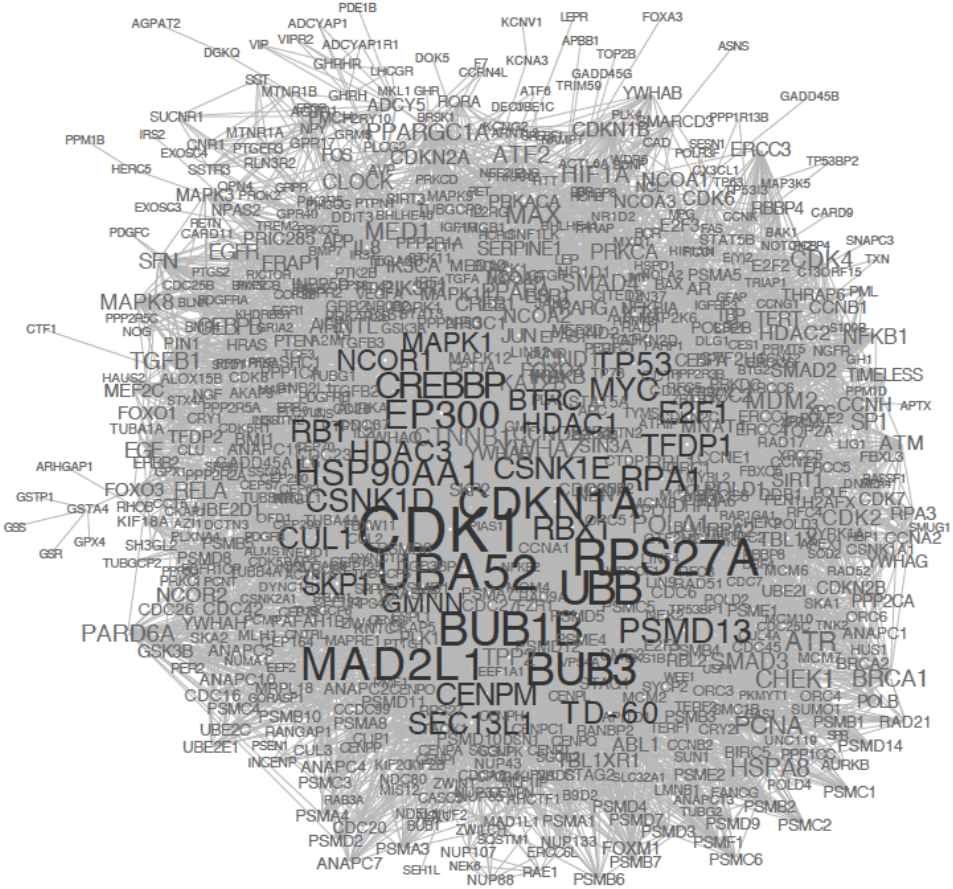
Multiscale Pathway Network. Nodes are gene symbols of identified molecular timekeepers found to interact with other timekeepers in STRING database for ‘databases’, evidence of interactions from curated pathway databases. Edges were included only if they linked two nodes that represented two or more temporal domains. Node size is proportional to node degree. Black indicates key drivers.

#### 3.1.4 Multiscale PPI Network

The vast majority of nodes with edges are part of a densely connected protein-protein interaction network (Fig. 6, Table S6), as observed above for the other nework types. This topological trait implies a large extent of regulatory communication across temporal domains. Node degree of the *p53* gene is substantial greater than the other key drivers, building on coexpression evidence of its prominent status in the multiscale temporal network. The highly connected gene *MDM2* fits this pattern in that it is regulated by *p53* but is also responsible for targeting *p53* for degredation.

**Figure 6.**
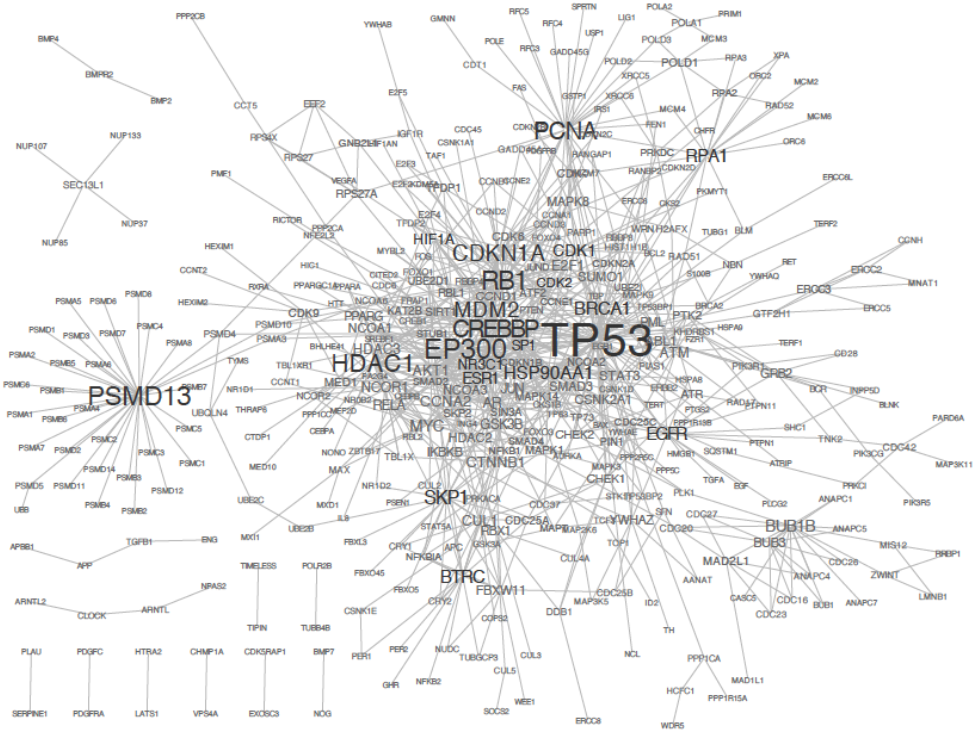
Multiscale PPI Network. Nodes are gene symbols of identified molecular timekeepers found to interact with other timekeepers in STRING database for ‘experiments’, experimental evidence of protein-protein interactions. Edges were included only if they linked two nodes that represented two or more temporal domains. Node size is proportional to node degree. Black indicates key drivers.

#### 3.1.5 Consensus Multiscale Network

The consensus network that included only edges appearing in greater than or equal to two network types (Fig. 7, Table S7), with 491 nodes and 812 edges, largely reflects aspects of the topologies observed for the single evidence types, but the similarities are greatest with evidence from PPI. Prominent within the most highly connected region are the genes *p53*, *EP300* and *CREBBP*, supporting the central importance of the latter two genes, identified across all three temporal scales (Table 1). A node that was highly connected in all the networks was *PCNA*, a circular ring-shaped protein that surrounds DNA, functioning as a molecular platform for replication and repair enzymes and involved in transcription, chromatin remodeling, chromatid cohesion, cell cycle regulation, and apoptosis [Cazzalini 2014]. *CREBBP* and *EP300* play a critical roll in genome stability by acetylation of PCNA, which causes its removal from DNA and degradation.

**Figure 7.**
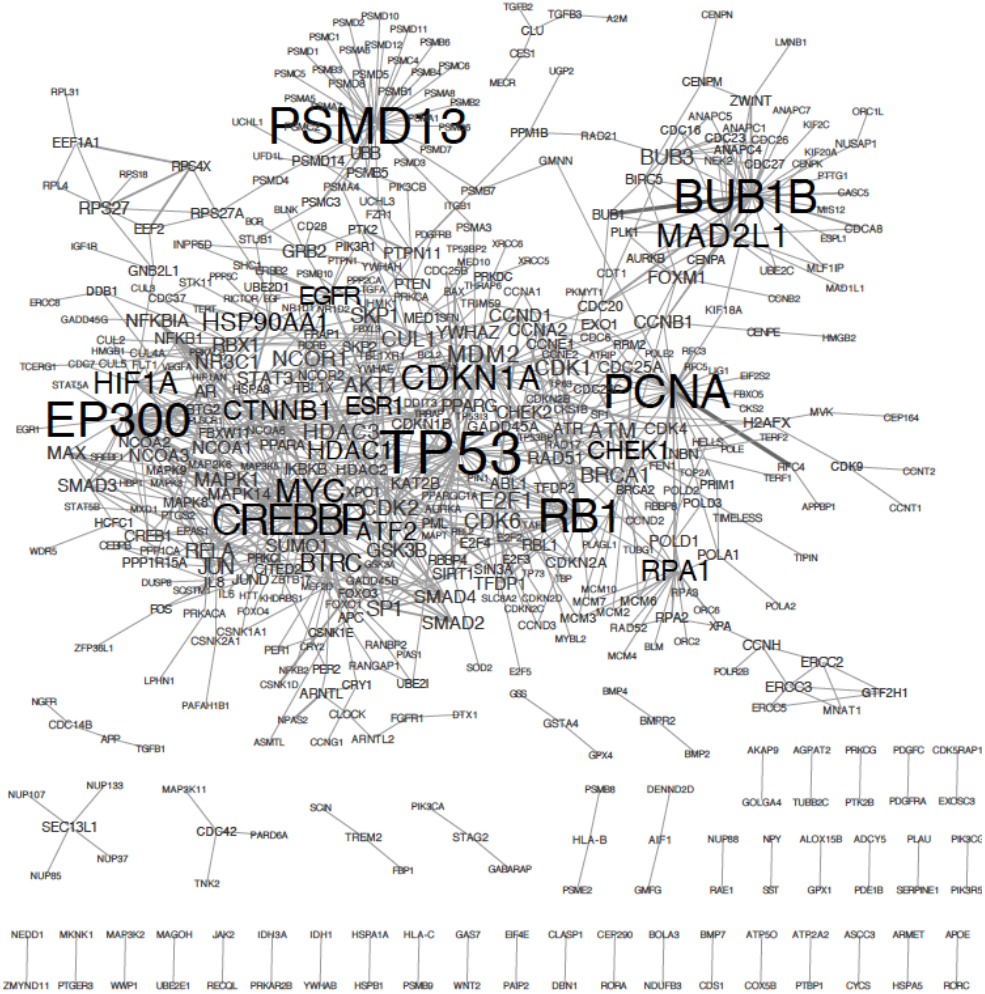
Consensus Multiscale Network. Edges were included only if they were in two or more multiscale networks described above. Node size is proportional to node degree, number of connections to the node. Black indicates key drivers. Edge width indicates the number of multiscale networks that include the edge, with the range being from 2 to 4.

To further investigate molecular functions of genes in the consensus network, PANTHER overrepresentation tests were conducted with Bonferroni corrections using the GO Ontology database (Mi, et al., 2016). After cell cycle related categories, which were enriched by design, categories relating to metabolic processes were most significantly enriched, for example, regulation of metabolic process (GO:0019222) (P = 6.07e-127). Also very highly enriched were the categories, cellular response to stress (GO:0033554), regulation of signal transduction (GO:0009966), regulation of cell communication (GO:0010646), and cellular response to stimulus (GO:0051716), (P = 1.56e-93, P = 1.82e-82, P = 5.90e-79, and P = 1.60e-78, respectively).

The most highly connected node by a large margin was *p53*, a gene that has been demonstrated to be involved in regulating a bewildering array of cell behaviors such as proliferation, senescence, cell death, growth, DNA repair, differentiation, stem cell reprograming, metabolism, and motility (Kruiswijk, et al., 2015), as well as mediating circadian regulation of cellular pathways (Gotoh, et al., 2014). *p53* fits the general pattern for nodes in the multiscale network in that it plays an important role in cellular signaling, responding to intrinsic or extrinsic signals to institute a transcriptional program to achieve different outcomes in a cell (Levine, et al., 2006).

## 4 Discussion

Genes in the multiscale network are likely to affect temporal processes not yet identified in the literature. Drugs, compounds, and exposures that affect transcription or function of genes in the network, or the regulatory relationships between those genes, may also affect multiple temporal biological processes, including processes that are beyond the scope of this analysis, such as growth and development or age at menarche. For example, given the important role of protein kinases in the consensus multiscale network, one might hypothesize that deficiency in Mg^2+^, an essential cofactor in many biological processes involving protein kinases, would result in dysregulation of temporal biological processes. In fact, it has been found that hypomagnesaemia can result in ventricular tachycardia, ventricular fibrillation, aging, young gestational age, tremors, and tumors (Long and Romani, 2014; Shah, et al., 2014). Despite substantial differences between multiscale network types, important reoccurring themes are apparent, such as signal transduction and *p53* pathways.

The consensus network is not intended to be comprehensive but rather a core set of genes and regulatory relationships that control human temporal biological processes across time scales. The fact that each edge was required to be supported by two types of evidence, each at a stringent confidence threshold, lends a high degree of certainty to the network. While it is possible that a small proportion of edges represent non-regulatory relationships, there is strong justification for interpreting the majority of edges as representing true interactions. Though at least three time scales are represented in the network and the majority of nodes in the network are densely connected, it is still not known definitively whether any particular node can be generalized to operate beyond its currently annotated temporal scales. We hope that these results will not only lead to studies that test hypotheses involving these genes individually but also to studies that explore new hypotheses about the role of the network in temporal biology.

## Conclusions

These results demonstrate a concept that has recently been advanced piecemeal by researchers in related disciplines, that is, the intertwined nature of the cell cycle, circadian, aging, metabolic, and signaling molecular processes. Further, the results highlight the possibility that a core network plays an important role in temporal biological processes generally. It also suggests the central role that signaling molecules play in temporal processes generally by integrating multiple signal inputs to institute a coordinated response.

